# Generalized peakgroup scoring boosts identification rates and accuracy in mass spectrometry based discovery proteomics

**DOI:** 10.1101/2022.11.03.515031

**Authors:** Aaron M. Scott, Christofer Karlsson, Tirthankar Mohanty, Suvi T. Vaara, Adam Linder, Johan Malmström, Lars Malmström

## Abstract

The statistical validation of peptide and protein identifications in mass spectrometry proteomics is a critical step in the analytical workflow. This is particularly important in discovery experiments to ensure only confident identifications are accumulated for downstream analysis and biomarker consideration. However, the inherent nature of discovery proteomics experiments leads to scenarios where the search space will inflate substantially due to the increased number of potential proteins that are being queried in each sample. In these cases, issues will begin to arise when the machine learning algorithms that are trained on an experiment specific basis cannot accurately distinguish between correct and incorrect identifications and will struggle to accurately control the false discovery rate. Here, we propose an alternative validation algorithm trained on a curated external data set of 2.8 million extracted peakgroups that leverages advanced machine learning techniques to create a generalizable peakgroup scoring (GPS) method for data independent acquisition (DIA) mass spectrometry. By breaking the reliance on the experimental data at hand and instead training on a curated external dataset, GPS can confidently control the false discovery rate while increasing the number of identifications and providing more accurate quantification in different search space scenarios. To first test the performance of GPS in a standard experimental environment and to provide a benchmark against other methods, a novel spike-in data set with known varying concentrations was analyzed. When compared to existing methods GPS increased the nunmber of identifications by 5-18% and was able to provide more accurate quantification by increasing the number of ratio validated identifications by 24-74%. To evaluate GPS in a larger search space, a novel data set of 141 blood plasma samples from patients developing acute kidney injury after sepsis was searched with a human tissue spectral library (10000+ proteins). Using GPS, we were able to provide a 207-377% increase in the number of candidate differentially abundant proteins compared to the existing methods while maintaining competitive numbers of global identifications. Finally, using an optimized human tissue library and workflow we were able to identify 1205 proteins from the 141 plasma samples and increase the number of candidate differentially abundant proteins by 70.87%. With the addition of machine learning aided differential expression, we were able to identify potential new biomarkers for stratifying subphenotypes of acute kidney injury in sepsis. These findings suggest that by using a generalized model such as GPS in tandem with a massive scale spectral library it is possible to expand the boundaries of discovery experiments in DIA proteomics. GPS is open source and freely available on github at (https://github.com/InfectionMedicineProteomics/gscore).

## 1 Introduction

One disadvantage of data independent acquisition (DIA) proteomics is that a spectral library based on previously identified peptides in a sample is required to interpret the complex signal and quantify peptides. In the past, this has made the practice of discovery proteomics in DIA difficult, as samples would first need to be run with data dependent acquisition (DDA) and extensive offline fractionation to reach the depth needed to see the benefits of the more sensitive DIA methods. However, recent computational advances in the development of fragment spectra prediction [19, 65, 59], and the creation of repository scale spectral libraries for selected organisms and sample types [51, 66, 39, 47, 6, 31] have allowed for the exploration of DIA as a means for discovery proteomics. Although these large libraries can help facilitate the identification of novel markers in a DIA experiment, they present significant computational difficulties, particularly when attempting to control the false discovery rate (FDR) using the target-decoy approach [14]. The increased search space of massive libraries can cause a decrease in sensitivity and statistical power as more false positives are introduced into the library, leading to less precursors being correctly identified when they are in fact in the sample [42, 18, 17]. In these cases, validation algorithms struggle to distinguish the true signal from the false, resulting in low numbers of validated peakgroups and imprecise FDR control. Attempts have been made to filter down these massive libraries to manageable sample-type specific libraries in a data-dependent fashion to a more manageable search space that statistical validation algorithms can readily deal with [26, 18]. However, if the library filtering is done too strictly potential true peptides are eliminated unnecessarily from the library and the benefits that could be gleaned from using these massive libraries are reduced, decreasing the potential depth of a discovery analysis. If the filtering is done too liberally, where too many false peptides are left in the library, the same issue arises where validation algorithms have trouble distinguishing true signal in the large search space. In both cases, this can result in an unreliable estimation of the FDR, so care must be taken when filtering large spectral libraries for analysis.

Alternative to the filtering and sub-setting of spectral libraries to analyze data with large spectral libraries, one case that has not been investigated with the same vigor is the effect of the choice of statistical validation algorithms on their ability to navigate this large search space and control the FDR in a stable manner. The algorithm of choice in the validation of DIA mass spectrometry data is the mProphet algorithm [49], which is similar to the percolator algorithm commonly used in DDA proteomics [58]. This mProphet algorithm, implemented in the python package PyProphet [50] for the use of validating DIA data extracted with the OpenSwath software [52], uses a semi-supervised method to combine all calculated sub-scores into a final classification score used to control the FDR. This method works by selecting positive training targets above a particular q-value cutoff in an iterative fashion once they are identified and validated by the algorithm in the previous iteration. The initial targets are identified using a selected sub-score and a broad q-value cutoff (typically 0.15 or 15% FDR) and the method progresses until no more new identifications are found to pass below a selected FDR threshold (1%). In most cases, where the library very closely matches with the data contained in the sample of interest, this method works exceedingly well. However, when searching a sample with a repository scale library, where the majority of precursors in the library are likely not contained in the sample, a situation arises where the normally “true” target labels are noisy. This means that the peakgroups labeled as a “target” in the spectral library and assumed to be in the sample are not actually contained in the sample. This is a particularly tricky area of machine learning research, and research has been done to develop methods that identify noisy labels and stabilize the training of models in the presence of noisy labels [43, 11]. PyProphet, and additionally the tool Percolator [58] designed for validating identifications in data dependent acquisition, attempt to mitigate this issue with the iterative semi-supervised learning approach, but when the number of true positives in the sample is so low, it is not guaranteed that each iteration can actually select true positives for training. If some true targets are identified, it would be an exceedingly small amount compared to the negative decoys in the data, creating a overwhelming class imbalance, which can destabilize the training of machine learning algorithms if not dealt with in an appropriate manner [2].

Instead of dealing with these types of noisy label and class imbalance problems on the fly during sample specific model training, we propose to create a stable, generalizable machine learning model that can be used to combine sub-scores from DIA extraction software by training on a curated data set of known true and false positive peakgroups. This approach has been demonstrated to work using static models with percolator in the context of DDA proteomics [58] but the static models are trained on the sample types that they are used to evaluate, so it is unclear if they would generalize to diverse datasets of unrelated sample types. We hypothesize that a good peakgroup is a good peakgroup, no matter the sample type, and that statistical validation models can be trained on an unrelated external data set if the data is curated properly. To that end, we have trained a generalizable scoring model and implemented a suite of algorithms to provide the stable validation of extracted peakgroups through the spectral library size-agnostic Generalizable Peakgroup Scoring (GPS) framework. Here, we compare the GPS to two existing validation tools, PyProphet[50] and Percolator[58] and evaluate the performance on 3 different novel datasets using different spectral libraries and search spaces. We demonstrate the use of GPS in a practical setting by analyzing blood plasma samples from patients with septic acute kidney injury using the repository scale Pan Human Library (PHL) [51] and evaluate the possibility of using this type of framework for the discovery of potential disease markers. GPS is a freely available as an open source Python package and command line tool on github (https://github.com/InfectionMedicineProteomics/gscore).

## 2 Results

### 2.1 Generalizable model training

To enable the generalized scoring of peakgroups, we first aimed to train a machine learning model that can detect true peakgroups in a sample with high precision. 129 yeast samples were run with varying gradient lengths (30, 45, 60, 90, 120 minutes) and analyzed with OpenSWATH [52] to generate 1893804 target peakgroups and 1857563 decoy peakgroups using the ms2-level sub-scores calculated by OpenSWATH as features. We split the dataset so 80% remained for training and 20% was reserved for evaluating the performance the machine learning models. Because many of the target labels in the dataset are noisy (ie. they originate from false targets) we developed an algorithm to filter out noisy labels from the dataset that we will refer to as the denoising algorithm. The denoising algorithm trains an ensemble of weak learners using bagging [7] that vote on each peakgroup from a particular mass spectrometric sample. If every single classifier in the ensemble voted that the peakgroup was a target, then the peakgroup is kept as a target in the training set, otherwise it is removed. The overall workflow for this method is depicted in Figure 1B. This results in a filtered training set with 1278834 true targets, marking a 15.53% decrease in the number of targets used for training, along with 1486043 decoy peakgroups. In Figure 1A the UMAP [38] projection shows that the cluster of decoy datapoints (labeled 0.0) is contaminated with target labels, making the cluster appear almost purple. After denoising, the effects can be visualized in Figure 1C which depicts the purity of the projected components for the target and decoy data points in the training set. To evaluate the performance of the denoising algorithm in generating a classifier to identify only true peakgroups, we trained 2 classifiers, 1 based on the unfiltered training data, and the 2nd of the filtered training set, and analyzed a held-out subset of the unfiltered training data that we will refer to as the testset. Overall on this testset we observed an 11.66% increase in the precision using the filtered model, from 89.19% to 99.58%, and a 96.13% decrease in the false discovery rate (FDR), from 10.81% to 0.42%. In D-E of Figure 1 the confusion matrices display the overall classification numbers that were used to calculate these metrics, and the overall score distributions for each method. The true and false labels widened substantially and the portion of targets deemed false more accurately model the decoy distribution in the filtered model as seen in D-E of Figure 1.

**Figure 1:**
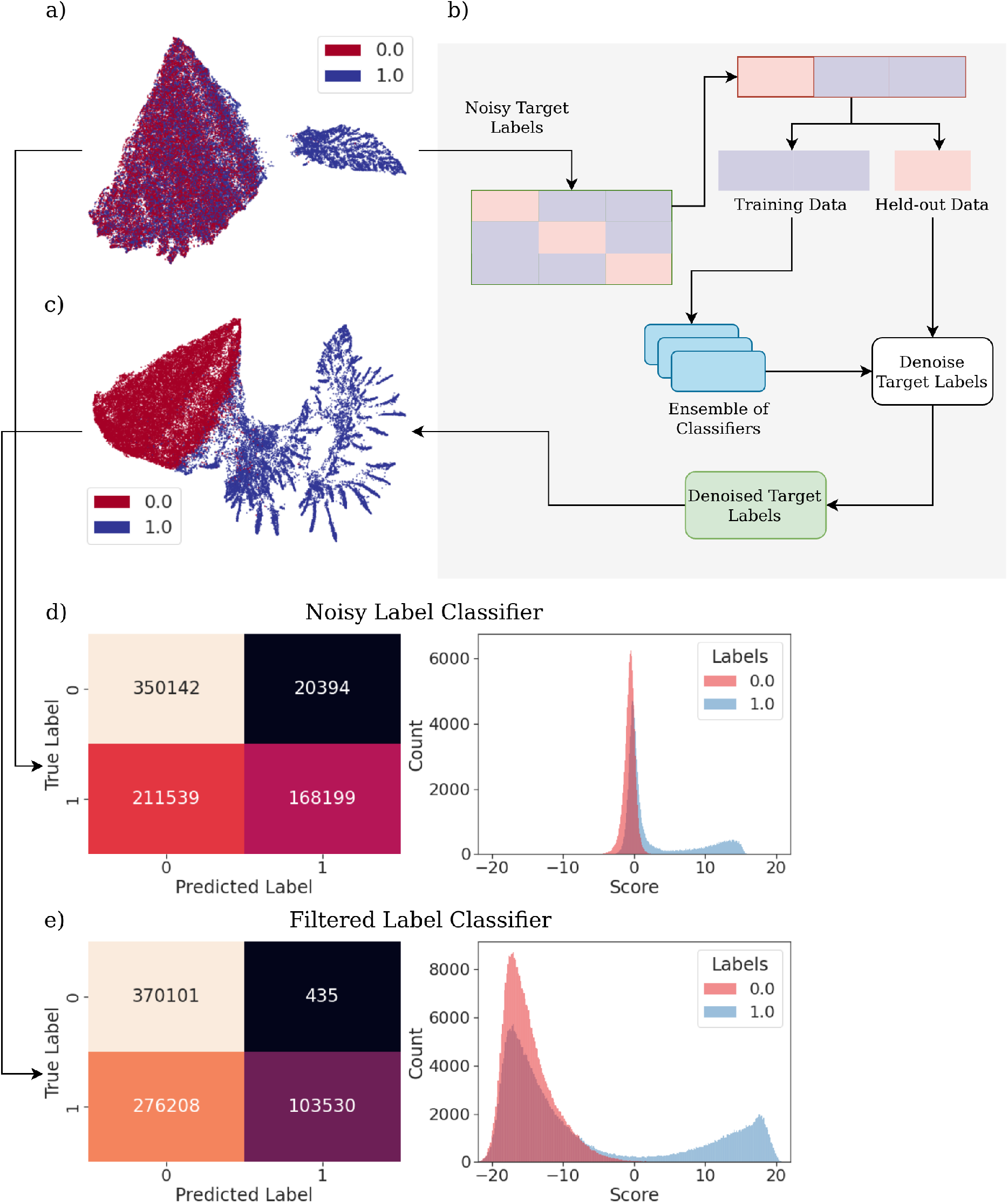
**a)** The UMAP projection of the unfiltered training labels. 1.0 is a target label and 0.0 is a decoy label. As visible by the projection, the cluster on the left corresponding to the decoy labels, is also fully populated with target labels. This means that during training, noisy target labels that are indistinguishable from decoy labels will be used as positive training instances when they should not be **b)** The overall workflow for denoising the noisy target labels in the training set. The training data is first split into k number of folds, for each held-out fold, the remaining training data is randomly sampled n-times with replacement to train n-classifiers that are then used to vote on the held-out data to determine if an instance is a true target or not. The probability threshold to consider a vote positive can be adjusted for stringency. **c)** The UMAP projection of the filtered training labels with a 0.75 probability vote cutoff after denoising. Each individual cluster is very pure, with only a few overlapping data points in the middle, meaning that the target labels used for training are much more confident that they are a true positive instance. **d)** The confusion matrices and score distributions for the classifier trained using the noisy unfiltered labels. **e)** The confusion matrices and score distributions for the classifier trained using the filtered labels.

### 2.2 Method Benchmark Comparison

To establish the validity of data produced using GPS with the filtered model described above, we first performed an analysis on samples with known ratios of 4X more yeast peptides in one sample group spiked-in into a constant mouse kidney proteome background. To verify that our method was validating correct peakgroups, we monitored the expected ratios to make sure that the quantification is accurate, while still maintaining high levels of identifications. We used OpenSwath [52] to perform signal extraction and then GPS, PyProphet [50], and Percolator [58] to validate the extracted peakgroups and compare between the different tools. To ensure that we were directly comparing the ability of only each machine learning tool to validate peakgroups, the scoring functions of each tool were used and then the scored samples were compiled and exported into global quantification matrices in the exact same way. At a 1% FDR, we observed 15.69% increase in precursors, an 18.20% percent increase in peptides, and a 14.79% increase in proteins identified by GPS compared to PyProphet and a 12.68% increase in precursors, a 14.40% increase in peptides, and a 4.7% increase in proteins identified by GPS compared to Percolator (Figure 2). To measure the accuracy of each validation method at identifying precursors correctly, we measured the number of true ratio validated identifications in a ±0.2 log2 fold-change (log2FC) window around the known yeast and mouse log2 fold concentrations as done previously [41, 26]. Compared to PyProphet, we observed a 24.30% increase in the number of ratio validated yeast identifications and a 28.83% increase in the number of mouse ratio validated identifications (Figure 2). Compared to Percolator, we observed a 68.52% increase in the number of ratio validated yeast identifications and a 74.06% increase in the number of mouse ratio validated identifications (Figure 2). The log2 fold change distributions and the ratio validation windows of interest for each method can be seen in A, C, and E of Figure 2. To measure the quantification accuracy beyond an arbitrary window of *±*0.2, we measured the average precision of ratio validations at increasing thresholds from the known concentrations of yeast and mouse proteins (Figure 2F). The average precision is calculated as the number of yeast identifications in the yeast window plus the number of mouse identifications in the mouse window divided by the total number of identifications found both windows at each given threshold. As seen in Figure 2F, GPS starts out with the highest precision closest to the known ratios and then drops as the windows widen while maintaining comparable precision to PyProphet until around the ±0.4 window when it is slightly lower than PyProphet (0.9744 PyProphet compared to 0.9746 GPS at the final cutoff). Percolator displays a lower precision at all windows compared to GPS and PyProphet, although it is worth noting that the precision of each method here is very high with 0.97 being the minimum precision found among all three methods.

**Figure 2:**
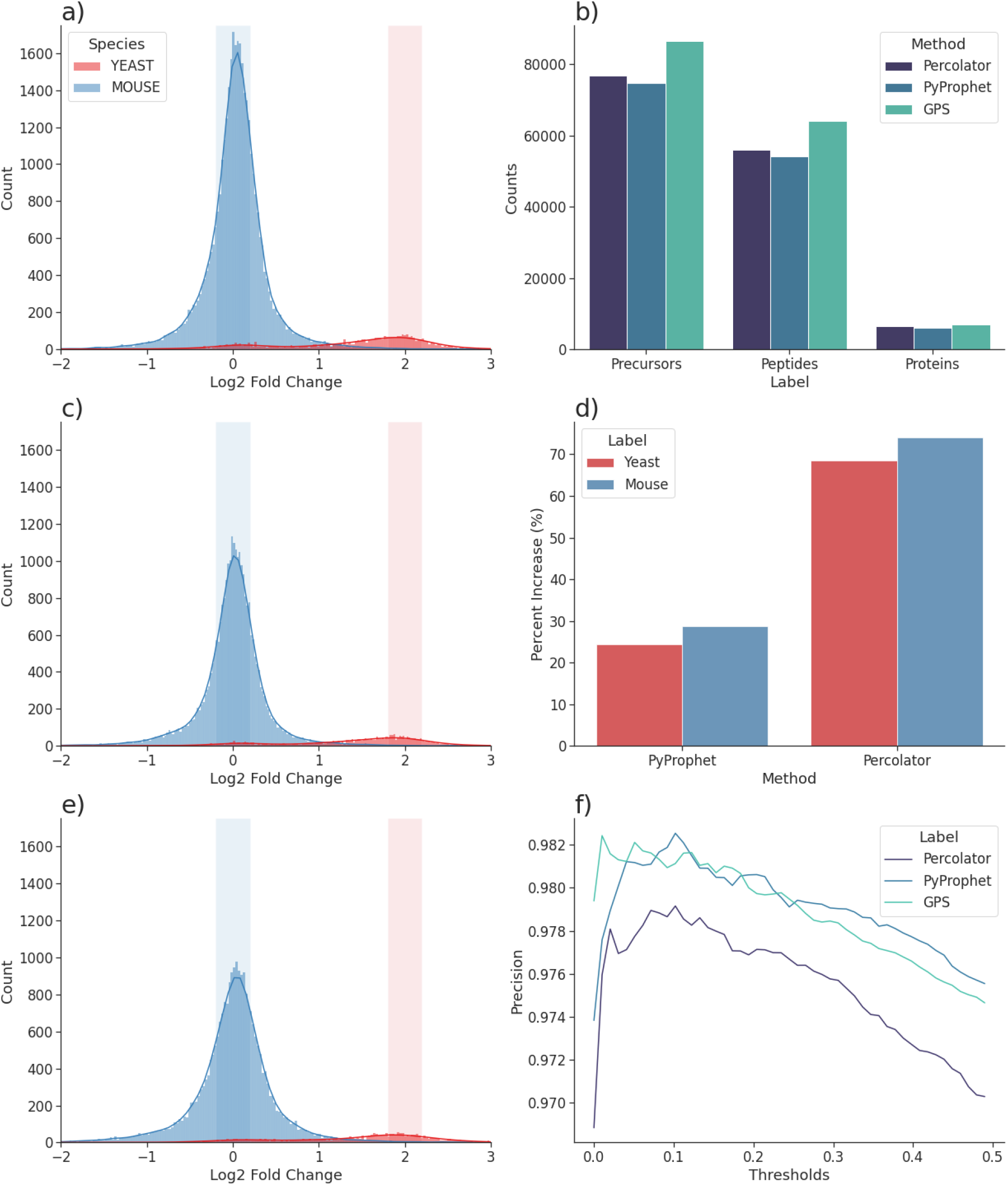
**a)** Distributions of the fold changes for GPS of the precursors with at least 2 measurements per group. The 0.2 log2FC windows around the expected mouse and yeast ratios are highlighted. **b)** Global precursor, peptide, and protein counts for each tool in the benchmark at a 1.0% global protein and peptide FDR. **c)** Distributions of the fold changes for PyProphet of the precursors with at least 2 measurements per group. The 0.2 log2FC windows around the expected mouse and yeast ratios are highlighted. **d)** Barplot showing the percent increase of ratio validated identifications for PyProphet and Percolator for the expected yeast ratio and the expected mouse ratio. This measures the accuracy and number of identifications in the indicated areas from a, c, and e in the figure. **e)** Distributions of the fold changes for Percolator of the precursors with at least 2 measurements per group. The 0.2 log2FC windows around the expected mouse and yeast ratios are highlighted. **f)** Lineplot that visualizes the precision of measurements of each tool at increasing log2FC threshold windows around the expected mouse and yeast ratios. Precision is measured by the proportion of correct identifications divided by all identifications within the tolerance windows.

### 2.3 Percentage of Incorrect Target (PIT) Analysis

To further establish confidence in the GPS method and to investigate the necessity of down-weighting decoys with the PIT during q-value calculation, we analyzed samples comprised of mouse kidney tissue and queried it with the same spectral library from the mouse-yeast spike-in analysis described above. The PIT (or pi0) correction has been used previously to allow for more identifications to be identified at the same FDR threshold due to the downweighting of decoys [27]. This method is implemented in both Percolator and PyProphet in order to increase the number of identifications overall, but it is possible that the inclusion of these identifications can lead to less accurate quantification and the inclusion of incorrect identifications in the resulting quantitative matrices. To test this, we used the denoising algorithm developed for model training above and applied it to the classification of putative peakgroups in a sample. The denoising voting ensemble proves accurate in identifying which portion of the target distribution should be classified as the false target distribution, and the overlay of these predicted distributions can be seeing in Figure 3A. To measure the true FDR, we used these false target classifications to perform a PIT correction during q-value calculation and then measured the number of yeast identifications that passed at the equivalent estimated FDR (Figure 3B). At a 1.0% FDR, GPS obtained the lowest Yeast FDR (1.94%), while the Yeast FDR calculated using GPS with the estimated PIT correction (2.15%) was 10.97% higher. The PyProphet (2.38%) and Percolator (2.13%) Yeast FDRs were 22.40% and 9.97% higher respectively. We also compared the true positive rate (TPR), measured using mouse identifications, and the false positive rate (FPR), measured using yeast identifications, at increasing FDR thresholds (Figure3C-D). At all thresholds, Percolator displays the lowest FPR, but also identifies significantly less mouse identifications at the same cutoffs (GPS with PIT estimation increases the number of identifications at a 1.0% FDR by 71.01% compared to Percolator). At a 1.0% FDR GPS with PIT estimation identifies the most correct mouse identifications, while maintaining a comparable FPR and Yeast FDR.

**Figure 3:**
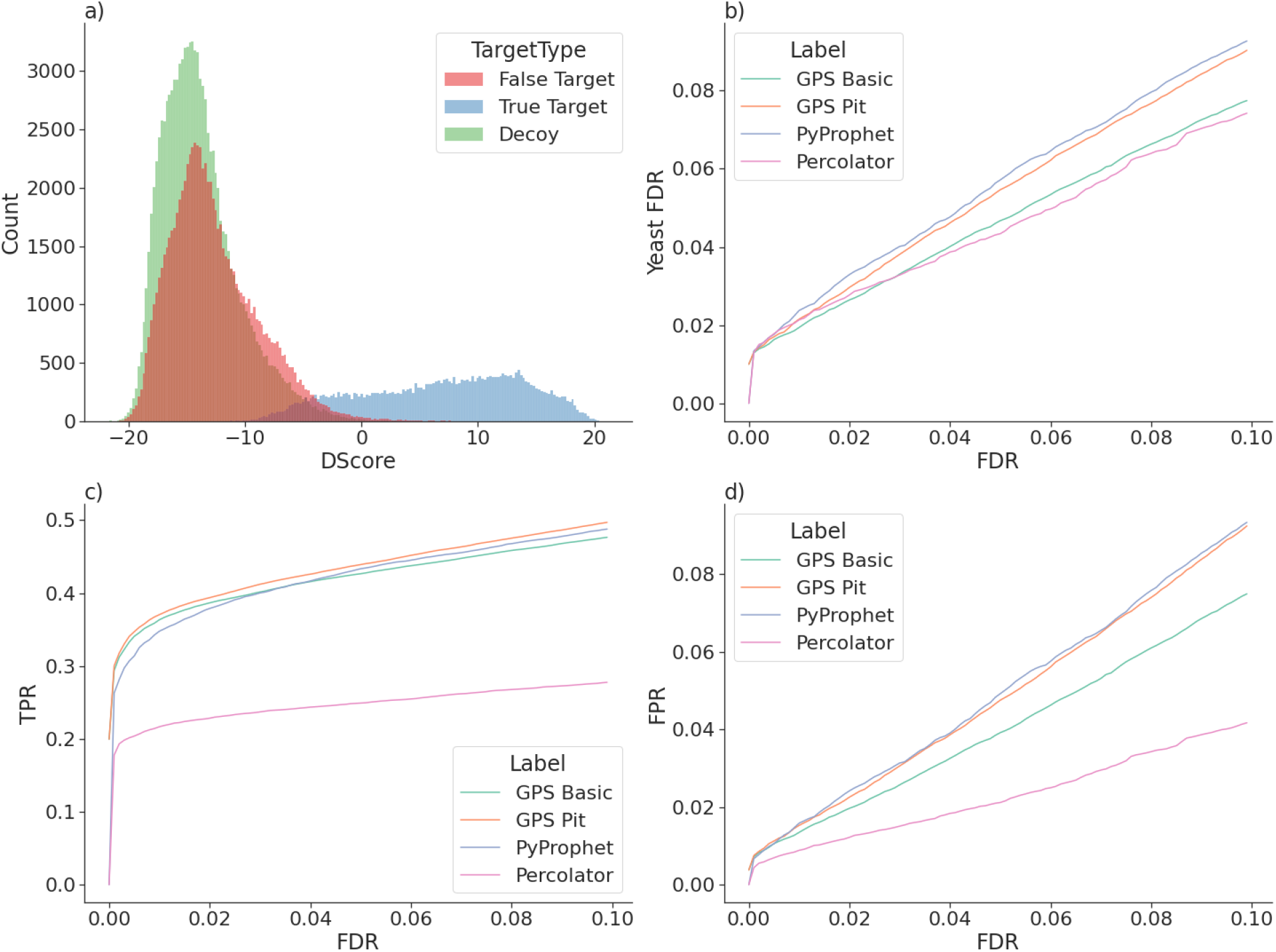
**a)** Density plots of the target, decoy, and false target distributions from a particular mouse kidney sample. The predicted false target distribution is determined using the denoising algorithm to correct q-value calculation during PIT estimation. The overlay of the decoy and false target distributions show the accuracy of the denoising method in predicting targets and false targets. **b)** The number of yeast identifications that pass at particular FDR thresholds for each measured method. Since no yeast identifications should be found in the sample, a completely linear relationship (y=x) would indicate perfect entrapment FDR control **c)** The true positive rate of mouse proteins identified with increasing q-value cutoffs for all measured methods. This acts as a proxy for the number of identifications at particular q-value cutoffs. **d)** The false positive rate as a measurement of the yeast peptides that pass at certain FDR thresholds compared to all yeast peptides in the library.

### 2.4 Acute kidney injury (AKI) analysis in patient blood plasma

To provide a biological context to the improvements provided by GPS, we analyzed 141 previously unpublished blood plasma samples from a subcohort of sepsis patients with acute kidney injury from the FINNAKI study [48, 20]. These 141 samples are comprised of 2 established subphenotypes, based on severity of the illness, that were developed from a combination of multiple clinical markers [61]. We interrogated this data using the repository scale Pan Human Library (PHL) (10838 proteins) [51] and an optimized PHL where we corrected the retention time and appended spectra from identifications obtained by searching the DIA data directly with MSFragger-DIA (v3.5) [30] [57] (10952 proteins). This analysis puts into context the benefits that GPS provides when querying a large search space and the benefit of using extensive curated repository spectral libraries in discovery DIA approaches and how they can be optimized.

#### 2.4.1 Large Search Space Comparison

When repository scale spectral libraries are used to query samples, classic peakgroup scoring methods can struggle to correctly validate a large portion of the proteins that truly exist in the sample due to the increased complexity of the data extracted. We analyzed the 141 blood plasma samples with the PHL and then used GPS, PyProphet, and Percolator to score the extracted signal, calculate q-values at the precursor, global peptide, and global protein levels, and aggregate the data into a quantitative matrix. The precursors validated by the 3 tools were rolled up into their proteins using a Python implementation of the relative quantification iq algorithm [46]. The resulting volcano plots in Figure 4A-C visualize the differentially abundant proteins between the 2 subphenotypes of AKI (81 samples were from subphenotype 2 and 60 samples were from subphenotype 1) above an absolute value of 1 log2FC and an adjusted p-value of 0.1. At these cutoffs GPS provides a 377.78% increase in the number of differentially expressed proteins compared to PyProphet, and a 207.14% increase in differentially expressed proteins compared to Percolator (Figure 4F). Percolator identified the most proteins at a global 1.0% FDR at the protein level with 937, compared to 768 using GPS, and 164 with PyProphet (Figure 4D). The quantitative matrices produced by Percolator and GPS contain 87.91% and 78.29% missing values (MV) respectively while PyProphet produced substantially less MVs overall (35.74%), although the number of proteins at the global level was also much lower (Figure 4). Overall GPS was able to identify 768 proteins globally, 91 differentially abundant proteins (adjusted p-value ¡ 0.1), and 381 potentially differentially abundant proteins (proteins found in at least 2 samples per group). The large increases in the number of differentially abundant proteins in Figure 4F that pass the specified 1.0 log2FC and 0.1 adjusted p-value cutoffs can be attributed to the fact that GPS provides a more complete data matrix in the sense that a greater number of proteins have multiple measurements per subphenotype, so more proteins can be accurately compared (a 38.55% increase compared to Percolator and a 154.0% increase compared to PyProphet).

**Figure 4:**
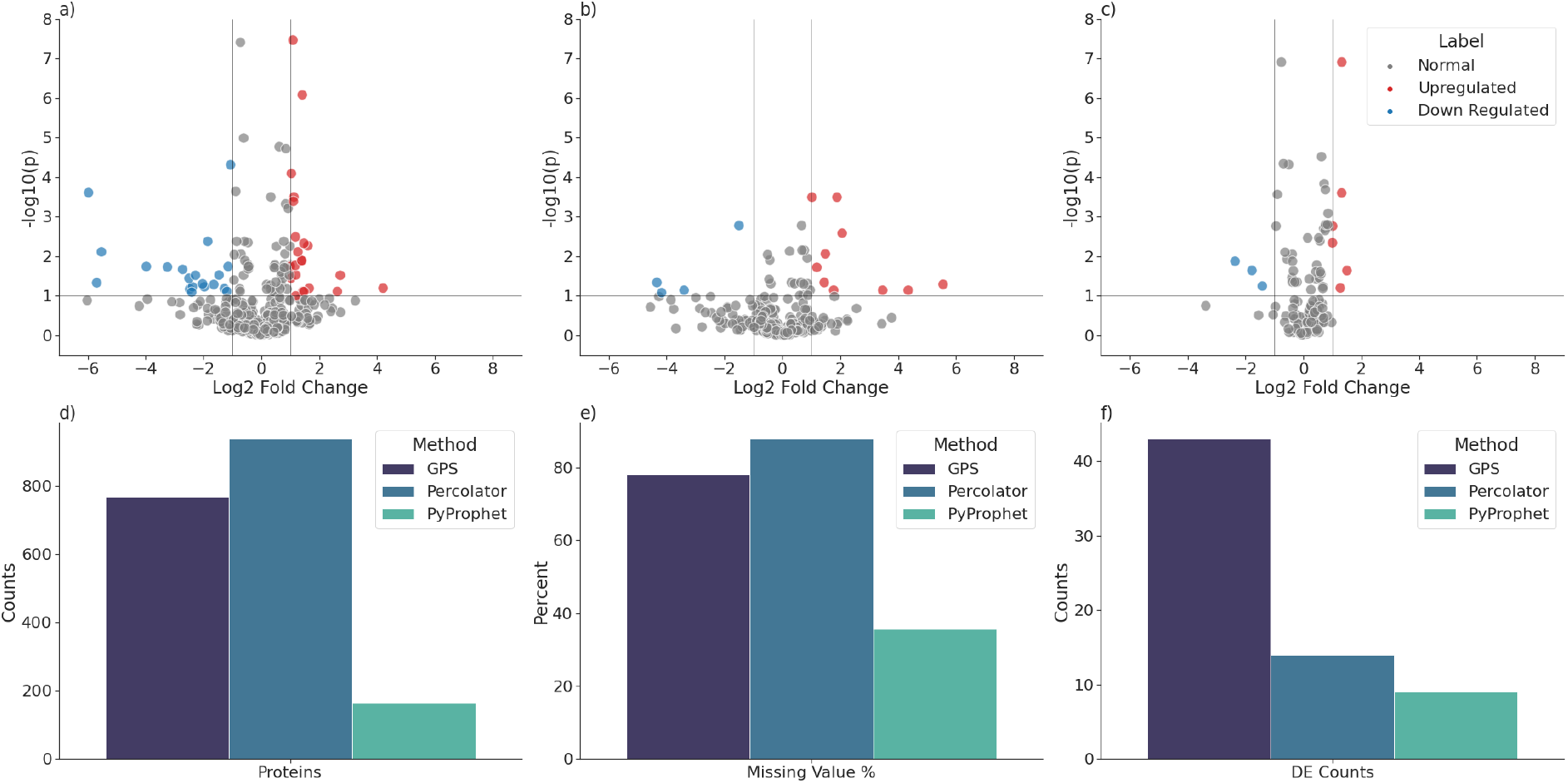
**a)** Volcano plot visualizing the distribution of differential abundance values (log2FC) for GPS on the AKI data and their -log10(adjusted p-value). The cutoffs are set at a 0.1 adjusted p-value and 1 log2FC. **b)** Volcano plot visualizing the distribution of differential abundance values (log2FC) for Percolator on the AKI data and their -log10(adjusted p-value). The cutoffs are set at a 0.1 adjusted p-value and 1 log2FC. **c)** Volcano plot visualizing the distribution of differential abundance values (log2FC) for PyProphet on the AKI data and their -log10(adjusted p-value). The cutoffs are set at a 0.1 adjusted p-value and 1 log2FC. **d)** Barplot of the overall global protein counts that pass a 1.0% FDR cutoff for the compared methods. **e)** Barplot of the percent of missing values in the dataset for proteins that pass a 1.0% FDR cutoff for the compared methods. **f)** Barplot of the number of differentially abundant proteins that pass a 1.0% FDR cutoff for the compared methods.

#### 2.4.2 Discovery DIA in plasma proteomics

To provide biological context, and to show the benefit of using full-tissue libraries in discovery DIA proteomics, we reanalyzed the 141 blood plasma samples using the optimized PHL with an iterative retention time alignment workflow described in the methods. Differential expression was performed between the 2 subphenotypes to identify potential proteins that could act as differentiating markers in stratifying AKI subtypes in sepsis. Using this optimized PHL and workflow we were able to detect 1205 proteins overall (a 56.90% increase compared to the standard PHL), 170 differentially abundant proteins (adjusted p-value ¡ 0.1), and 651 potentially differentially abundant proteins (proteins found in at least 2 samples per group). In Figure 5A we can see the overall distribution of differentially abundant proteins in the dataset.

**Figure 5:**
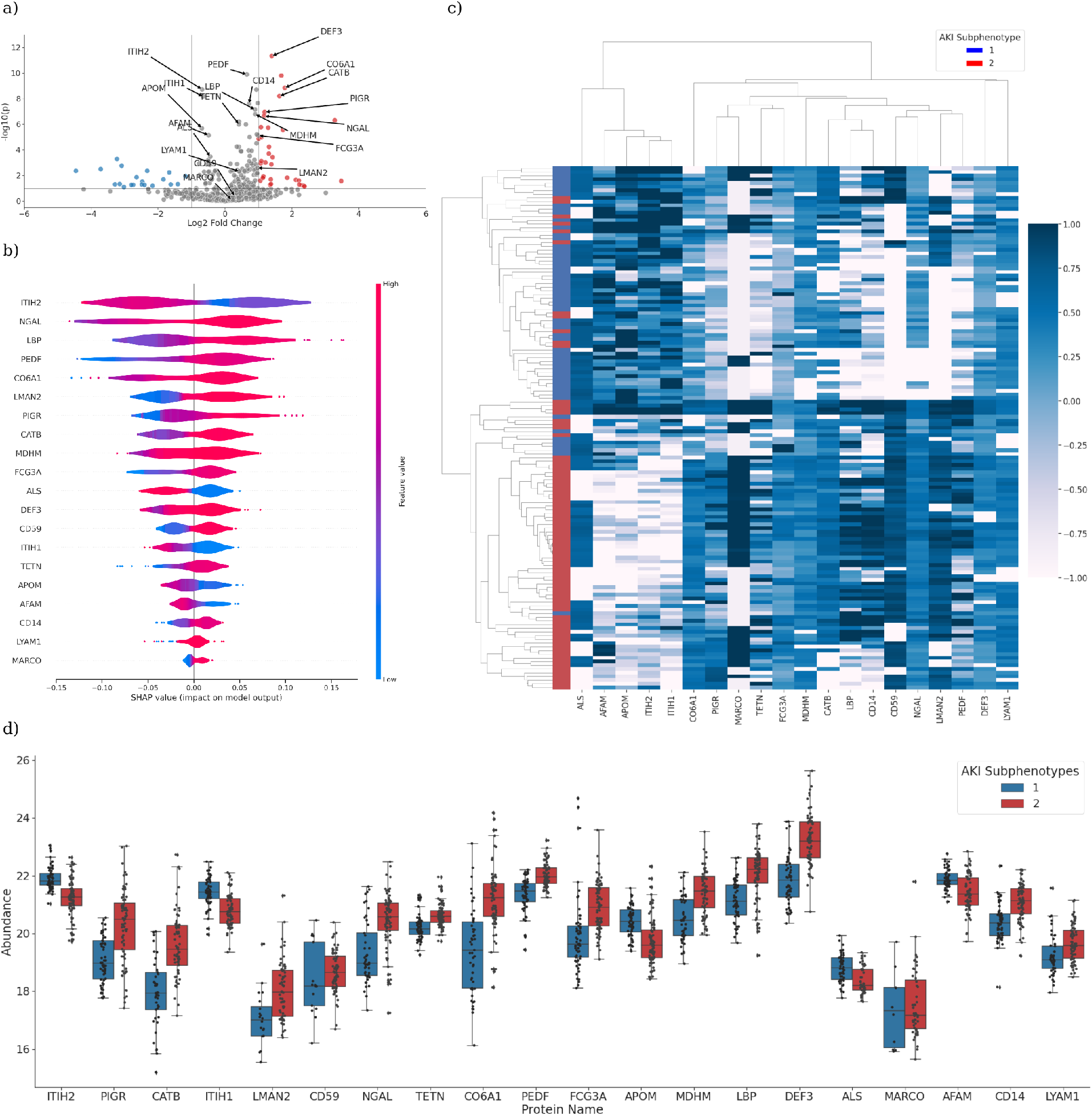
**a-c)** Volcano plots for the differential abundant proteins analyzed with the optimized PHL library. Red corresponds to proteins upregulated in subphenotype 2 and blue corresponds to proteins upregulated in subphenotype 1. The labeled proteins correspond to the top 20 most important proteins found that differentiate between subphenotypes with a Random Forest model. It is worth noting that many of the most important proteins do not fall outside of the arbitrary cutoffs commonly used to identify proteins of interest in biomarker studies. **b)** Violin plots of the SHAP scores for the top 20 proteins indicated in Figure5A. A red color indicates a higher abundance for a protein and a blue color indicates a lower abundance for a protein. SHAP scores greater than 0.0 on the x-axis indicate features that drive classification towards subphenotype 2 and scores less than 0.0 indicate protein values that drive classification towards subphenotype 1. **c)** Heatmap and cluster analysis of the top 20 proteins identified using SHAP scores from the Random Forest model. There is a clear grouping of the subphenotypes (0.76 accuracy), and 2 main clusters of proteins, the first indicating proteins of lower abundance in subphenotype 2 and the second of higher abundance in subphenotype 2. **d)** Box plots of the abundance of the top 20 selected proteins compared between subphenotypes. It is interesting to see that many of the proteins do not have clear separation of the means between the groups, but are still distinctly important in classifying between subphenotypes.

#### 2.4.3 Machine learning aided differential expression

Utilizing the boosted number of potential marker proteins detected with the optimized PHL, we wanted to investigate the potential to train a machine learning model to differentiate between the 2 subphenotypes of AKI mentioned above, interpret the importance of each protein in the model, and use this interpretation to guide the selection of interesting proteins for further study. Using 10-fold cross validation we trained a Random Forest classifier which resulted in a mean accuracy of 0.84 (0.12 standard deviation). To quantitatively understand the importance of each protein used in the model as a feature we leveraged the explainable machine intelligence algorithms in the SHAP python package[35]. From the trained model, we calculated the SHAP values for each protein and have visualized the top 20 proteins with the greatest mean importance (Figure 5B). Using these top 20 proteins we performed an average hierarchical clustering with correlation similarity using a maximum of 2 groups and saw that the subphenotypes clustered together with 76% accuracy (0.76 Rand score) and we could easily visualize a clear separation of the 2 groups (Figure 5C). For the purposes of the visualization, we capped the normalized protein abundances at -1.0 and 1.0. Additionally, we annotated the volcano plot in Figure 5A to showcase that the top 20 most important proteins in differentiating between the 2 subtypes using the trained random forest model would not pass the arbitrary canonical cutoffs used to identify interesting differentially expressed proteins. In Figure 5e we plot these top 20 selected proteins to visualize their abundances across the 2 subphenotypes.

## 3 Discussion

With the implementation of our new methods, we show an increase in the precision of quantification, while also increasing the number of identifications when compared to the established methods. We were also able to demonstrate how these established methods can suffer when the spectral library search space is too large and does not match the sample, while GPS is able to score data in this scenario in a stable manner. These combined improvements allow for the deep and in-depth analysis of plasma proteome samples using repository scale spectral libraries to boost the power of discovery DIA experiments.

### 3.1 Benchmark comparison with existing methods

PyProphet and Percolator both use a semi-supervised learning technique to train on a particular data set by identifying new true targets every iteration and re-training until the number of peakgroups that pass a q-value threshold no longer increase. This type of method can provide great results on individual experiments, particularly by increasing the number of identifications, as it maximizes the local number of targets that are validated at the end, but could potentially introduce false positives that are elucidated only when analyzing the precision of the quantitative accuracy on known spiked-in proteins. It has also been observed that these semi-supervised methods can struggle when the number of samples analyzed is relatively low (low-n) [17] or when the spectral library used contains substantially more proteins than it is possible to find in the sample (Figure 4). To minimize these issues, we looked to leverage one of the core strengths of machine learning algorithms by training a generalizable model on a diverse dataset that can accurately separate true targets from false targets independent of the search space size. Since it is difficult to directly quantify the performance of new computational methods based on individual metrics, we created a species mixture data set of known ratios of yeast peptides spiked in to a constant mouse kidney proteome background. This allows us to evaluate the efficiency of different peakgroup validation methods and compare them directly based on the number of identifications and the accuracy of quantification based on how close the produced quantitative values are to their expected ratios. We demonstrate here that the optimized GPS model outperforms existing methods based on the number of global identifications because the performance of GPS is not hindered by the specificity of the samples being analyzed since we trained the generalizable model on millions of extracted peakgroups from an external dataset. We also demonstrate a boost in accuracy and precision of quantification due to the automated curation of the training data to ensure only true peakgroups are included when building the model without over-fitting on experiment specific data. These performance improvements suggest that generalized scoring models can be leveraged in diverse experiment types to provide boost identification numbers and quantitative accuracy without the need to train new models for each experiment. Additionally, there is no need to optimize hyperparameters to squeeze the best performance out of GPS, as the generalized model will score peakgroups accurately in a stable manner no matter the conditions of the data. Existing methods can be optimized in different conditions, but a meticulous optimization of hyperparameters is required. This non-trivial time consuming parameter optimization task can be avoided completely by implementing peakgroup scoring with generalizable models such as with GPS.

### 3.2 PIT estimation

PIT estimation has become an established method to boost the number of identifications that pass through at a given FDR cutoff in mass spectrometry proteomics. However, predicting this percentage of false targets is computationally difficult, as it is unknown which targets are in fact false, or where to split the target distribution to estimate pi0. Existing methods use certain heuristics to estimate pi0 by calculating the difference between the decoy counts and target counts at certain score cutoffs [27], or provide naive counts based on the number of targets below a 1.0% FDR, but this can lead to inaccurate results depending on the shapes of the score distributions. To accurately estimate pi0, we leveraged the ensemble denoising algorithm used to filter the training data for the high precision GPS model. By measuring the number of targets below a 100% vote percentage we can accurately model the false target distribution and use it to down-weight the decoy counts when calculating q-values. This can considerably increase the number of identifications that pass at certain thresholds, especially in large search space scenarios (ie. respository scale spectral library searches). It is possible to underestimate or overestimate pi0 by changing the probability thresholds of the denoising classifier, and these can be leveraged in different situations depending on the goal of the particular experiment. The implications of false target prediction goes beyond the downweighting of decoys during q-value calculation, it also opens up the possibility of decoy free scoring methods that utilize the predicted false target distributions directly to calculate q-values. This could massively cut down the search space, particularly in discovery DIA settings using repository scale or deep learning predicted libraries.

### 3.3 Boosted performance in library-sample size mismatches

In theory, it is particularly beneficial to search blood plasma samples with massive scale complex libraries to delve deeper into the proteome as it could be possible to pick up and identify potential disease markers missed in standard analysis, especially in the case of low abundant proteins. In practice, however, classic peakgroup validation algorithms struggle when the true signal extracted with the library is much smaller than the library itself, or when the library size is significantly larger than the amount of proteins contained in the sample. For example, when a blood plasma sample is searched with a repository scale library containing 500,000 precursors there may only be 15,000 identifiable precursors in the sample, so a massive imbalance in search space is created. As seen in Figure 4, the existing semi-supervised algorithms can struggle in these search space imbalances and lead to significantly less candidate differentially expressed proteins being identified. One potential reason for this is that the library search space imbalance creates a situation where the true target labels are extremely noisy, ie. only 3% of the target peakgroups extracted by the library are true peakgroups. Semi-supervised methods attempt to get around this in an iterative fashion using an algorithm where new targets are selected based on q-value cutoffs [50, 58]. When the true target to decoy ratio is so small, it is not guaranteed that a correct set of true targets are picked up for training during the semi-supervised iteration, creating a massive imbalance in the ratio of true training targets to false training targets. There are a few different ways to try and mitigate training set imbalances, such as down-sampling the majority class, up-sampling the minority class [2], which Percolator does, or by using the synthetic minority oversampling technique (SMOTE) where synthetic training instances are created based on the distributions of features found in the minority class [15]. It is also possible to provide the class ratios to certain machine learning algorithms to ensure that over-represented classes do not dominate the training loops. Unfortunately, these methods are not implemented in all common peakgroup validation algorithms, so even though some may work effectively in situations where the library size is very similar to the sample composition, they can still struggle in situations as the library grows larger and larger compared to the sample. In the case of GPS, instead of choosing to implement imbalanced learning methods to validate the data with sample specific classifiers we chose to build a static classifier that leverages modern machine learning technologies and generalizes to diverse sample and experiment types. In these cases, it is also extremely beneficial that there is no need to optimize the parameters of GPS to get these results, it can realize these improvements out of the box.

### 3.4 Analysis of AKI samples in human plasma

Taking the algorithmic benefits of GPS into consideration, we decided to apply the new method to the computationally difficult data set of AKI plasma samples described above to provide some biological context. Using our 1205 globally identified proteins, and instead of selecting candidate proteins for further analysis based on arbitrary q-value and log2 fold change cut offs, we trained a machine learning model to differentiate between the 2 subphenotypes of AKI using the candidate differential proteins in the dataset and then calculated the importance of each protein to the model during classification using SHAP values [35]. The high accuracy and separating power of this panel of 20 proteins indicates that they would provide a good starting point for investigation as potential clinical markers. In fact, many of these proteins have already been identified and studied as potential clinical sepsis markers or markers for infection and inflammation [21, 44, 60, 16, 45, 36, 37, 67, 64, 10, 62, 23, 9, 55, 1, 25]. The majority of the top 20 proteins were up-regulated in the more severe subphenotype 2. Some examples which were higher in subphenotype 2 are the neutrophil-enriched NGAL (Neutrophil-gelatinase associated lipocalin), a previously described marker of severe AKI [60]; CD14 (Cluster of differentiation 14), a marker of monocyte infiltration; serum acute phase protein LBP (Lipopolysaccharide binding protein); neutrophil-specific DEF3 (Neutrophil defensin 3); and mitochondria specific MDHM (Malate dehydrogenase, mitochondrial). Proteins that were downregulated in subphenotype were the liver-specific acute phase protein ITIH-1, -2 (Inter-alpha-trypsin inhibitor heavy chain-1, -2)[56] and the albumin-like AFAM (Afamin) whose levels are known to be lowered with increase in sepsis severity [33]. This panel could further be expanded to any protein that has a significant weight in classifying the severity of AKI based on a combination of SHAP values and differential expression analysis in an effort to identify novel disease biomarkers.

These findings suggest that it is possible to identify potential sepsis markers in plasma samples and accurately quantify them using repository scale spectral libraries and statistical validation with GPS. These added benefits could significantly aid in the stratification of sepsis subphenotypes by allowing for a deeper exploratory investigation of the plasma proteome on a systematic basis and the informed data-driven selection of potential biomarkers for further validation. This approach would also generalize to other biological conditions or diseases easily, providing a systematic method towards discovery DIA for a wide range of experiment types.

## 4 Conclusion

We have proposed GPS as a method for the statistical validation of DIA mass spectrometry data, and provided evidence that generalized scoring models can outperform dynamically trained models especially in a large search space environment. GPS provides stable validation that leads to more accurate downstream quantification and provides evidence that sophisticated generalized scoring models can be used in tandem with massive scale spectral libraries to support the development of discovery proteomics in DIA mass spectrometry.

## 5 Methods

### 5.1 Datasets

| Dataset | MS Acquisition | N Samples | Technical Replicates | Usage |
| --- | --- | --- | --- | --- |
| HFX-SPD | DIA | 129 | All | Generalizable model training |
| Mouse-Yeast | DDA | 60 | 10 | Spectral library generation |
| Mouse-Yeast | DIA | 60 | 10 | GPS Benchmark |
| Mouse Kidney | DIA | 31 | None | Entrapment FDR Analysis |
| AKI | DIA | 141 | None | Large Search Space Comparison<br>Optimized Library Analysis<br>ML Differential Expression |

#### 5.1.1 GPS model training data: HFX-SPD

In order to provide a chromatographically diverse set of training data, we used a data set comprised of 129 different samples of 500ng Yeast tryptic digest (Promega) with varying gradient lengths (30, 45, 60, 90, 120 minutes) and acquired with DIA. This data set will be referred to as the HFX-SPD data set.

#### 5.1.2 Spike-in data

We generated a spike-in dataset of known concentrations of Yeast tryptic digest peptides (Promega) spiked in to a constant mouse kidney background. The mouse kidney material was obtained from the a previous project [40]. All animal use and procedures were approved by the local Malmö/Lund Institutional Animal Care and Use Committee, ethical permit number 03681-2019. C57BL/6 mice (Janvier, Le Genest-Saint-Isle, France) were sacrificed, and kidneys were isolated into a tube containing DPBS and silica beads (1 mm diameter, Techtum). The kidneys were then homogenized using MagNAlyser (Roche) and stored at -80°C. Homogenates were then thawed and centrifuged at 10,000g for 10’ at 4°C. The supernatant containing the soluble proteins was collected and protein content was estimated using BCA (Pierce). 25ug of protein was taken for reduction, alkylation, digestion and C18 clean up, as described below. A serial dilution series of yeast peptides (1X, 2X, 4X, 8X, 16X, 32X) was performed, and 10 technical replicates of each concentration were sampled for a total of 60 samples. Internal retention time peptides were also added in to each sample. Each of the 60 samples were analyzed on the an Orbitrap HF-X using both DDA and DIA.

#### 5.1.3 Mouse kidney data

31 mouse kidney samples were selected from a previous study [40] to provide a base for the entrapment FDR analysis. The samples are from the same study as the mouse kidney material used to prepare the spike-in data, and follow the same ethical considerations and approvals (ethical permit number 03681-2019), as well as sample preparation methods described above.

#### 5.1.4 AKI data

AKI plasma samples used in the study belong to the FINNAKI study [48], a prospective, observational, multicenter study evaluating development of AKI in ICU patients with sepsis and septic shock. AKI was defined according to the Kidney Disease: Improving Global Outcomes (KDIGO) criteria based on changes in serum creatinine[29]. 51% of the patients developed AKI within the first 5 days in the ICU, with 30% diagnosed *<*12 hours from admission. Approximately 100 patients each developed stage 1, 2, and 3 AKI. 91 patients received RRT and the 90-day mortality for AKI patients was 33.7%. 141 samples were chosen for up to 5 time points from 23 acute kidney injury patients. The patients were from 2 distinct subphenotypes that were previously defined using a panel of clinical markers and latent class analysis [61].

### 5.2 Mass spectrometry sample preparation and data acquisition

#### 5.2.1 AKI sample preparation and analysis

141 samples were chosen for up to 5 time points from 23 acute kidney injury patients. All sample preparation steps, including desalting and protein digestion, used the Agilent AssayMAP Bravo Platform (Agilent Technologies, Inc.) per manufacture’s protocol. Each plasma sample was diluted 1:10 (100-mM ammonium bicarbonate (AmBic); Sigma-Aldrich Co, St Louis, MO, USA), and 10 l of each diluted plasma sample were transferred to a 96-well plate (Greiner G650201) where 40 μL of 4 M urea (Sigma-Aldrich) in 100 mM AmBic was manually added with a pipette for a final volume of 50 μL. The proteins were reduced with 10 μL of 60 mM dithiothreitol (DTT, final concentration of 10 mM, Sigma-Aldrich) for one hour at 37 °C followed by alkylation with 20 μL of 80 mM iodoacetamide (IAA, final concentration of 20 mM, Sigma-Aldrich) for 30 min in a dark at room temperature. The plasma samples were digested with 2 μg Lys-C (FUJIFILM Wako Chemicals U.S.A. Corporation) for five hours at room temperature and further digested with 2μg trypsin (Sequencing Grade Modified, Promega, Madison, WI, USA) overnight at room temperature [5]. The digestion was stopped by pipetting 20 μL of 10% trifluoroacetic acid (TFA, Sigma-Aldrich) and the digested peptides were desalted on Bravo platform. To prime and equilibrate the AssayMAP C18 cartridges (Agilent, PN: 5190-6532), 90% acetonitrile (ACN, Sigma-Aldrich) with 0.1% TFA and 0.1% TFA were used, respectively. The samples were loaded into the cartridges at the flow rate of 5 μL/min. The cartridges were washed with 0.1% TFA before the peptides were eluted with 80% ACN/0.1% TFA. The eluted peptides were dried in a SpeedVac (Concentrator plus Eppendorf) and resuspended in 25 μL of 2% ACN/0.1% TFA. The peptide concentration was measured using the Pierce Quantitative Colorimetric Peptide Assay (Thermo Fisher Scientific, Rockford, IL, USA). The samples, 10 μL, were diluted with 10 μL 2% ACN/0.1% TFA and spiked with synthetic iRT peptides (JPT Peptide Technologies, GmbH, Berlin, Germany) before liquid chromatography-mass spectrometry (LC-MS/MS) analysis.

#### 5.2.2 All other sample preparations and analysis

All protein samples were denatured with 8M urea and reduced with 5 mM Tris(2-carboxyethyl)phosphine hydrochloride, pH 7.0 for 45 min at 37 °C, and alkylated with 25 mM iodoacetamide (Sigma) for 30 min followed by dilution with 100 mM ammonium bicarbonate to a final urea concentration below 1.5 M. Proteins were digested by incubation with trypsin (1/100, w/w, Sequencing Grade Modified Trypsin, Porcine; Promega) for at least 9 h at 37 °C. Digestion was stopped using 5% trifluoracetic acid (Sigma) to pH 2 to 3. The peptides were cleaned up by C18 reversed-phase spin columns as per the manufacturer’s instructions (Silica C18 300 Å Columns; Harvard Apparatus). Solvents were removed using a vacuum concentrator (Genevac, miVac) and were resuspended in 50 μl HPLC-water (Fisher Chemical) with 2% acetonitrile and 0.2% formic acid (Sigma).

Peptide analyses were performed on a Q Exactive HF-X mass spectrometer (Thermo Fisher Scientific) connected to an EASY-nLC 1200 ultra-HPLC system (Thermo Fisher Scientific). Peptides were trapped on precolumn (PepMap100 C18 3 μl; 75 μl X 2 cm; Thermo Fisher Scientific) and separated on an EASY-Spray column (ES903, column temperature 45 °C; Thermo Fisher Scientific). Equilibrations of columns and sample loading were performed per manufacturer’s guidelines. Mobile phases of solvent A (0.1% formic acid), and solvent B (0.1% formic acid, 80% acetonitrile) was used to run a linear gradient from 5% to 38% over various gradient length times at a flow rate of 350 nl/min. The 44 variable windows DIA acquisition method is described by Bruderer et al [[8]]. MS raw data was stored and managed by openBIS (20.10.0) [[3]] and converted to centrioded indexed mzML files with ThermoRawFileParser (1.3.1) [[24]].

### 5.3 Spectral library creation

#### 5.3.1 Experiment specific spike-in library

An experiment specific library for the spike-in data was built by analyzing the samples acquired using DDA using FragPipe (v18.0). First the samples were searched using MSFragger (v3.5) [30] with default parameters using a fasta file of Swiss-prot reviewed saccharomyces cerevisiae and mus musculus proteomes concatenated with reverse sequence decoy proteins. Peptide spectrum matches (PSMs) were validated using Percolator[58]. The Philosopher toolkit (v4.4.0) was used to perform protein level FDR control with ProteinProphet, generate downstream reports, and filter the resulting identifications[34]. The python package easypqp was then used to convert and format the library for use by OpenSwath and the OpenMS (2.6.0) tool chain for formatting PQP files was used to optimize the assays in the library [54]. 10 spiked-in retention time peptides (iRT) were used for alignment and retention time correction for each sample.

#### 5.3.2 Pan Human Library

The Pan Human Library [51] was downloaded in its original form and then converted to a PQP spectral library using the OpenSwathAssayGenerator tool available from OpenMS [54].

#### 5.3.3 Optimized Pan Human Library

To augment the PHL with additional identifications and correct the retention time to the experiment at hand, we first searched the 141 AKI plasma samples using MSFragger-DIA (v3.5) [30, 57]. Using the resulting set of identifications, shared precursors between the PHL and direct search were selected for retention time alignment. LOWESS was first used to smooth the correlation between the direct search results and the PHL and then an interpolated univariate spline function was fit on top of this to adjust the retention time in the direct search to the scale of the PHL. The shared proteins between the 2 libraries were replaced in the PHL with proteins, and associated precursors, from the direct search, and the proteins not contained in the PHL were appended to the library. 10 spiked-in retention time peptides (iRT) were used for alignment and retention time correction for each sample.

### 5.4 Refined OpenSwath analysis

We adapted the previously published DIAnRT workflow [13] to optimize signal extraction at the sample level before combining the analysis to control for the global FDR. To do this, a first iteration is performed where sub-optimal retention time peptides are provided to align the experimental chromatograms to the spectral library. The first pass of signal extraction is scored and validated, and then the highest scoring peakgroups from a specified number of bins are selected and written out to a sample specific retention time library. This sample specific retention time library is used to align and correct the retention time to the spectral library with more stringent parameters in a second pass. Again, the second pass analysis is scored and validated and the highest scoring peakgroups for a specified number of bins are extracted and used as sample specific retention time libraries for a final pass. The final pass uses these sample specific retention time libraries to additionally correct mz and rentention time in an even more stringent manner. These final validated peakgroups are then used to infer peptide and proteins in a global manner to control for FDR and are finally combined in an output matrix that can be analyzed using the DPKS package mentioned above. Software to perform the sample specific retention time library extraction can be found in combination with the GPS python package and complete snakemake workflows and corresponding command line options for the different tools used can be found at (https://github.com/InfectionMedicineProteomics/gscore).

### 5.5 GPS

#### 5.5.1 Implementation

GPS is a Python library and command line utility for the generalized statistical validation of peakgroups. The source can be found here (https://github.com/InfectionMedicineProteomics/gscore). GPS leverages the package numpy [22] for efficient processing of numerical data, scikit-learn, sklearn and xgboost for implementing machine learning algorithms, and numba [32] for its just-in-time (JIT) compilation that compiles Python to machine code for optimization in performance critical areas of the library.

#### 5.5.2 Denoising algorithm

The denoising algorithm used to filter the HFX-SPD training set and predict the false target distribution for PIT correction is based on the concept of bagging from machine learning[7]. The data to be analyzed is first split into k number of folds (default is 10, and what is used throughout the study). Each fold is scored by training an ensemble of n logistic regression classifiers (default is 10, and what is used throughout the study) using stochastic gradient descent [53]) on data that is randomly sampled with replacement from the data left out of the selected k-fold. The ensemble of classifiers is then used to score the k-fold data, providing an average target probability for each peakgroup if the fold, and voting on each peakgroup to determine the vote percentage. A vote is considered a positive vote if the predicted probability for the individual classifier in the ensemble exceeds a threshold.

#### 5.5.3 Filtering training data

In an attempt to remove the noisy labels from the training dataset, the denoising algorithm described above was used to calculate a vote percentage for each peakgroup. If the calculated vote percentage was 100% then the peakgroup was kept as a true target. The probability to accept a positive vote was set at 0.75 to more strictly filter out potential false positives at the risk of losing some true identifications in the dataset. The negative training set, the decoy peakgroups, remained unfiltered.

#### 5.5.4 PIT Estimation

To estimate the PIT, the denoising algorithm was used to calculate the vote percentage for each peak-group in the dataset, where only the top scoring peakgroup was kept per precursor. For Figure3 0.75 was the probability threshold for a positive vote, and all peakgroups below a 100% vote percentage are considered false targets. The PIT is calculated by then dividing the number of false targets by the number of decoys.

For global PIT calculations on the peptide and protein level, the PIT is estimated by counting the number of peptides or proteins below a 1% FDR cutoff and then dividing that by the number of decoys.

#### 5.5.5 Peakgroup scoring and q-value calculation

The algorithm used to score each peakgroup in GPS is very straightforward. The peakgroups and their associated sub-scores are read in and parsed into a data structure that maps each scored peakgroup to the potential precursor. All of the peakgroups are scored using the filtered model trained above. Each peakgroup is additionally ran through the denoising algorithm to provide the needed scores to estimate the PIT. After scoring, the top ranked peakgroup for each precursor is selected to represent that precursor and the PIT is estimated. Q-values are calculated using an implementation of the qvality algorithm[28], where an interpolated spline is fit to the distributions of the target and decoy scores. A q-value for a particular peakgroup is calculated by first integrating the area under the curve of the decoy distribution from that particular score to the max and multiplying it by the PIT to get the decoy counts at a particular score threshold. The target counts are then obtained by integrating the area under the curve of the target distribution from that particular score to the max. Finally, the decoy counts are divided by the target counts plus the decoy counts to calculate a q-value, which can be used to filter for a given FDR. The highest scoring peakgroup for each precursor and the corresponding scores and q-value are written out to a file for downstream processing.

#### 5.5.6 Global FDR control

Global FDR control is implemented in a similar manner to PyProphet[50], where all scored samples in an experiment are aggregated and the highest scoring precursor is selected to represent either the peptide or the protein at the desired level. Once the highest scoring precursors are selected, q-values are estimated using the method described above and PIT corrected using the global PIT correction method. The resulting scoring models are exported as serialized Python objects that can then easily be used from the command line by GPS to export an annotated quantitative matrix.

#### 5.5.7 Quantitative matrix export

GPS can aggregate all scored samples, and the global peptide and protein models, into a quantitative matrix for downstream analysis. Each sample is read in to a data structure that filters the samples in the precursor based on their individual q-values. Once all samples have been parsed, they are annotated with their global peptide and protein level q-values using the score distribution objects that were previously built. The resulting annotated quantitative matrix is then written out for downstream analysis by the tool of your choosing.

### 5.6 Downstream statistical analysis

All down stream analysis was performed using the Data Processing Kitchen Sink (DPKS) Python package for general purpose data processing of mass spectrometry proteomics data. (https://github.com/InfectionMedicineProteomics/DPKS)

For all datasets, a retention time-mean sliding window normalization method was used based on the implementation in the NormalyzerDE R package [63]. Proteins were quantified for the AKI analysis using an implementation of the iQ relative quantification algorithm [46] (also known as MaxLFQ[12]). Differential expression was preformed using linear models, at the precursor level for the spike-in analysis and protein level for the AKI analysis. Multiple testing correction was performed using the Benjamini-Hochberg method [4].

All of these methods, including other options, are available in the DPKS package.

#### 5.6.1 Machine learning enhanced differential expression

In order to provide context and a ranking to the differentially expressed proteins, we trained a Random Forest Classifier (RFC) using the potentially differentially abundant proteins to classify between the subphenotypes in the AKI analysis. Missing values in the quantitative matrix were first imputed with zero values, as it is assumed if the protein was not quantified and identified that it is not in the sample. The protein quantities are then scaled to remove the mean and scale to unit variance. The model was evaluated using 10-fold cross validation to provide mean accuracy and confusion matrices. We used the SHAP[35] python package to then calculate the relative importance of each protein in differentiating between the subphenotypes of AKI. It was then possible to rank the differentially expressed proteins by their relative importance instead of setting arbitrary p-value and log2FC cutoffs to identify proteins for further investigation.

